# Association between schizophrenia and both loss of function and missense mutations in paralog conserved sites of voltage-gated sodium channels

**DOI:** 10.1101/246850

**Authors:** Elliott Rees, Noa Carrera, Joanne Morgan, Kirsty Hambridge, Valentina Escott-Price, Andrew J. Pocklington, Alexander L. Richards, Antonio F. Pardiñas, GROUP Investigators, Colm McDonald, Gary Donohoe, Derek W Morris, Elaine Kenny, Eric Kelleher, Michael Gill, Aiden Corvin, George Kirov, James T. R. Walters, Peter Holmans, Michael J. Owen, Michael C. O’Donovan

**Affiliations:** MRC Centre for Neuropsychiatric Genetics and Genomics, Division of Psychological Medicine and Clinical Neurosciences, School of Medicine, Cardiff University, Cardiff, UK; Centre for Neuroimaging and Cognitive Genomics (NICOG), National University of Ireland Galway, Galway, Ireland; Department of Psychiatry & Trinity Translational Medicine Institute, Trinity College Dublin, Dublin, Ireland

## Abstract

Sequencing studies have highlighted candidate sets of genes involved in schizophrenia, including activity-regulated cytoskeleton-associated protein (ARC) and N-methyl-d-aspartate receptor (NMDAR) complexes. Two genes, *SETD1A* and *RBM12*, have also been associated with robust statistical evidence. Larger samples and novel methods for identifying disease-associated missense variants are needed to reveal novel genes and biological mechanisms associated with schizophrenia. We sequenced 187 genes, selected for prior evidence of association with schizophrenia, in a new dataset of 5,207 cases and 4,991 controls. Included were members of ARC and NMDAR post-synaptic protein complexes, as well as voltage-gated sodium and calcium channels. We observed a significant case excess of rare (<0.1% in frequency) loss-of-function (LoF) mutations across all 187 genes (OR = 1.36; *P*_*corrected*_ = 0.0072) but no individual gene was associated with schizophrenia after correcting for multiple testing. We found novel evidence that LoF and missense variants at paralog conserved sites were enriched in sodium channels (OR = 1.26; *P* = 0.0035). Meta-analysis of our new data with published sequencing data (11,319 cases, 15,854 controls and 1,136 trios) supported and refined this association to sodium channel alpha subunits (*P* = 0.0029). Meta-analysis also confirmed association between schizophrenia and rare variants in ARC (*P* = 4.0 × 10^−4^) and NMDAR (*P* = 1.7 × 10^−5^) synaptic genes. No association was found between rare variants in calcium channels and schizophrenia.

In one of the largest sequencing studies of schizophrenia to date, we provide novel evidence that multiple voltage-gated sodium channels are involved in schizophrenia pathogenesis, and increase the evidence for association between rare variants in ARC and NMDAR post-synaptic complexes and schizophrenia. Larger samples are required to identify specific genes and variants driving these associations.

**Author Summary:** Common and rare genetic variations are known to play a substantial role in the development of schizophrenia. Recently, sequencing studies have started to highlight specific sets of genes that are enriched for rare variation in schizophrenia, such as the synaptic gene sets ARC and NMDAR, as well as voltage-gated sodium and calcium channels. To confirm the role of these gene sets in schizophrenia, and identify specific risk genes, we sequenced 187 genes in a new sample of 5,207 schizophrenia cases and 4,991 controls. We find an excess of protein truncating mutations with a frequency <0.1% in all 187 targeted genes, and provide novel evidence that mutations altering amino acids conserved across sodium channel proteins are risk factors for schizophrenia. Through meta-analysing our new data with previously published sequencing data sets, for a total of 11,319 cases, 15,854 controls and 1,136 trios, we increase the evidence for association between rare coding variants and schizophrenia in voltage-gated sodium channels, as well as in synaptic gene sets ARC and NMDAR. Although no individual gene was associated with schizophrenia, these findings suggest larger studies will identify the specific genes driving these associations.

## Introduction

Schizophrenia is a highly heritable polygenic disorder [1]. Collectively, common alleles contribute up to half of the genetic variance in schizophrenia liability [2, 3], and 145 distinct loci have currently been associated with the disorder at genome-wide levels of significance in the most recent genome-wide association study [4]. Schizophrenia risk is also conferred by rare mutations including copy number variants (CNVs) [5, 6] and rare coding variants (RCVs) [7, 8], each of which sometimes occur as *de novo* mutations [9, 10].

Studies of RCVs have the potential to inform schizophrenia pathogenesis since they can pinpoint specific functional variants in individual genes. However, to date, only two genes, *SETD1A* [11] and *RBM12* [12], have been strongly implicated. A major limiting factor, as for studies of common variants, is that for complex disorders, large samples are generally required to obtain robust results in case-control [13] studies. To date, the largest published sequencing studies of schizophrenia have involved around 5,000 cases, 9,000 controls and 1,000 parent-proband trios [7, 11], which are an order of magnitude smaller than recently published schizophrenia SNP genotyping studies of common risk alleles (e.g. 40,675 cases and 64,643 controls [4]). Nevertheless, whole exome sequencing studies have provided important clues to the pathophysiology of schizophrenia. For example, proband-parent trio based studies have shown *de novo* RCVs to be significantly enriched among glutamatergic post-synaptic proteins, in particular, the activity-regulated cytoskeleton-associated protein (ARC) and N-methyl-d-aspartate receptor (NMDAR) complexes [9]. These synaptic gene sets, first associated with schizophrenia through studies of *de novo* CNVs [10], have also shown evidence for association in independent case-control CNV [14] and sequencing datasets [7, 15]. More recently, in an extension of the Swedish sample used by Purcell *et al* 2014 [15], the authors documented an elevated exome-wide burden of ultra-rare, protein disruptive alleles, which was concentrated among 3,388 neuron-specific genes, particularly those that are expressed at synapses, including the ARC and NMDAR complexes [7]. Additionally, the enrichment of RCVs in schizophrenia has been shown to be concentrated among 3,488 genes that are depleted for loss-of-function (LoF) mutation in large population cohorts [16, 17].

In the current study, we performed targeted sequencing of 187 genes, selected for prior evidence for association with schizophrenia (Table S1), in 5,207 cases and 4,991 controls, none of which have contributed to previous schizophrenia sequencing studies. Among these targeted genes, we had complete membership of 4 gene sets, each of which has been postulated to be implicated in schizophrenia through rare variant analysis [7, 9, 10, 15, 18]; ARC and NMDAR post-synaptic protein complexes [9, 10], and voltage-gated sodium [18] and calcium channels [15]. Multiple voltage-gated calcium channels have also been strongly implicated through common variant studies [19]. Our primary aims were to a) investigate association between schizophrenia and RCVs in these gene sets, and b) identify individual genes that might drive the gene set associations. The remainder of the genes targeted for sequencing were selected on the basis of supportive evidence from at least two sources (see methods).

Most recent studies of RCVs in schizophrenia have focused on LoF alleles. However, it is clear that missense alleles also contribute to schizophrenia risk [7, 9], but in contrast to LoF alleles, *in silico* methods cannot distinguish at high sensitivity and specificity between missense alleles that alter the function of the encoded protein and those that are benign. Recently, it has been shown that restricting analyses to missense variants affecting amino-acids that are conserved within paralogous gene families improves power for identifying pathogenic alleles [20]. Given that two of our targeted gene sets consist of paralogous gene families (voltage-gated sodium and calcium channels), we exploited this approach to analyse missense variants that are more likely to have an adverse effect on protein function [20].

Finally, to maximise power, we combined the new sequencing data with independent, published schizophrenia case-control (Swedish [7] and UK10K [11] datasets) and trio exome-sequencing data (see methods), yielding a combined analysis of RCVs in a total of 11,319 cases, 15,854 controls and 1,136 trios.

## Results

### Mutation burden

In the targeted sequence sample, we performed six primary tests of mutation burden across all 187 targeted genes: LoF, nonsynonymous damaging and nonsynonymous variants, each under two allele frequency thresholds; < 0.1% and singletons. A significant (*P*_*corrected*_ < 0.05) excess of LoF mutations (< 0.1% in frequency) was observed in cases (Table 1), who had a mean excess of 0.013 LoF mutations per person across the 187 targeted genes (Supplementary Table S2). There was no significant difference in the rate of any other class of allele (Table 1). Although not part of our primary analysis, we note no difference between cases and controls in the rate of synonymous mutations (frequency <0.1%) (OR (95% CI) = 1.02 (0.94−1.08); *P* = 1), suggesting the enrichment of LoF mutations in cases is unlikely to be due to technical artefacts.

**Table 1.**
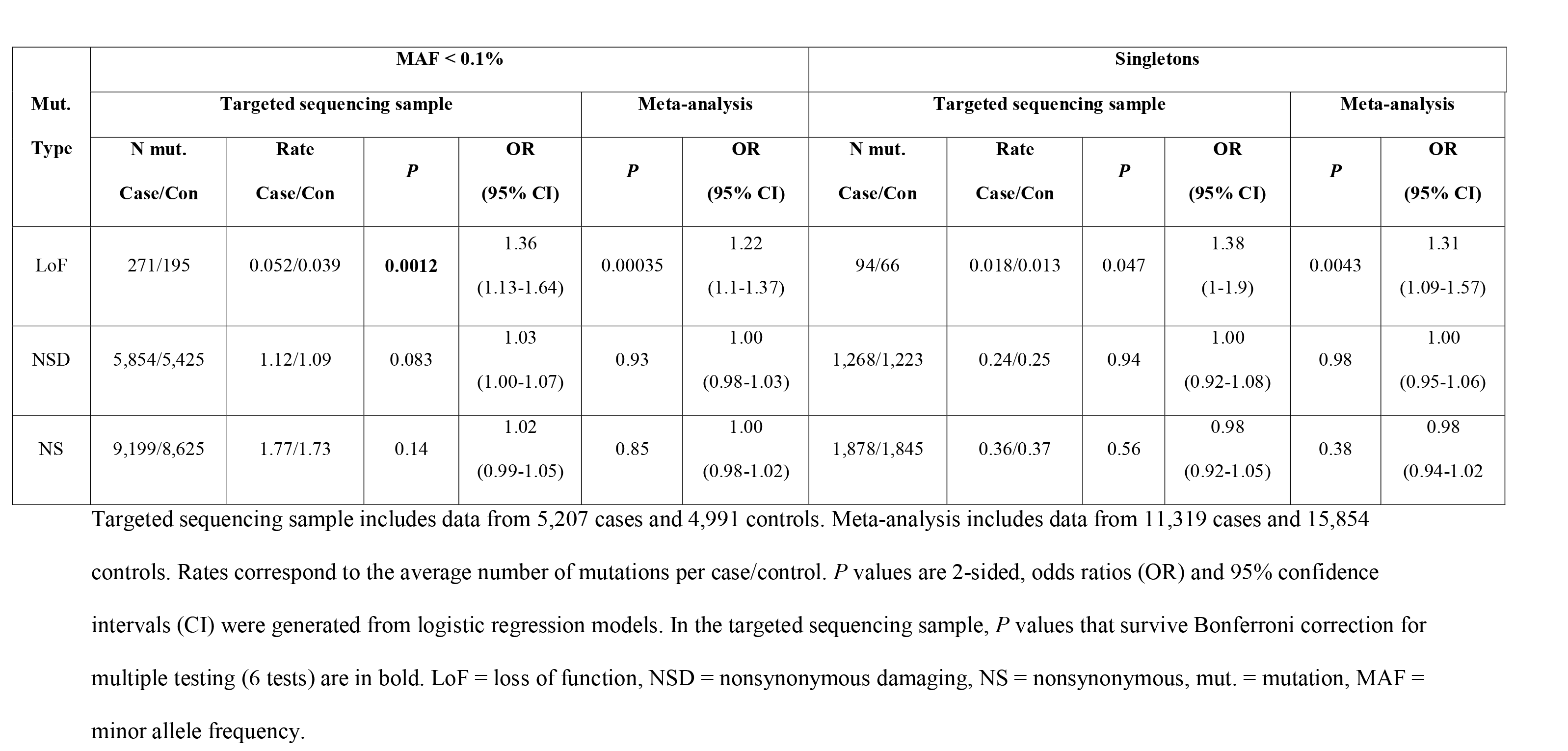
Mutation burden in 187 targeted gene

Meta-analysis with two previously published case-control exome sequencing datasets (Sweden and UK10K, see methods for detail) strengthened the evidence for an increase in LoF mutations (frequency <0.1%) in cases (Table 1 and Supplementary Table S3). The additional support came entirely from the Swedish rather than the UK10K dataset (Sweden OR (95% CI) = 1.27 (1.08−1.51; *P* = 4.8 × 10^−3^; UK10K OR (95% CI) = 0.95 (0.74−1.2); *P* = 0.66). The results contributing to the meta-analysis are presented in Supplementary Table S3.

We partitioned the 187 genes into those intolerant of LoF mutation (pLi scores > 0.9 in nonpsych-ExAC data [16]) and those that are not intolerant (pLi ≤ 0.9). Meta-analysis of the case-control data showed association between schizophrenia and rare (frequency <0.1%) LoF mutations was driven by LoF intolerant genes (106 genes with pLi > 0.9: OR (95% CI) = 1.63 (1.33 – 2.0); *P* = 2.9 × 10^−6^. 81 genes with pLi ≤ 0.9: OR (95% CI) = 1.09 (0.95 – 1.24); *P* = 0.21). The difference in effect size between pLi and non-pLi burden tests was significant (Z-test *P* = 0.0006).

### Gene set analysis

In the targeted sequencing analysis, as only LoF mutations with a frequency <0.1% were significantly enriched in cases after correcting for multiple testing, we tested this class of mutation for gene set enrichment.

*ARC and NMDAR:* In the targeted sequencing sample, cases had a higher rate of LoF mutations (frequency <0.1%) in ARC and NMDAR sets (Fig 1). When meta-analysed with published case-control datasets, we found strong evidence that LoF mutations in NMDAR genes were associated with schizophrenia (*P* = 1.6 × 10^−4^, Fig 1 and Supplementary Table S4), but weaker evidence for association with ARC genes (*P* = 0.047, Fig 1 and Supplementary Table S4).

**Fig 1.**
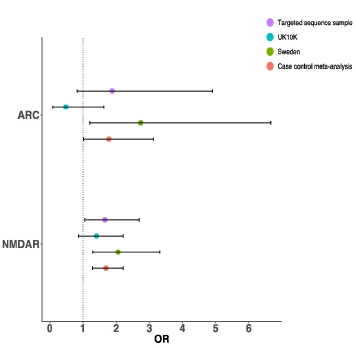
Case-control analysis of rare (frequency <0.1%) loss of function mutations insynaptic gene sets ARC and NMDAR.

To summarize the current status of RCVs in the above gene sets, we combined the case-control meta-analysis data with the *de novo* mutation data, selecting the class of *de novos* reported to be most strongly enriched in these gene sets (nonsynonymous *de novo* mutations in ARC and LoF *de novo* mutations NMDAR) in the previous work [9]. In the trio data, nonsynonymous and LoF *de novo* mutations were associated with ARC (*P* = 0.0015) and NMDAR (*P* = 0.014), respectively. Combining the *de novo* enrichment results with the case-control meta-analysis results (LoF, frequency < 0.1%), both ARC (*P* = 4.0 × 10^−4^) and NMDAR (*P* = 1.7 × 10^−5^) were associated with schizophrenia (Table 2).

The ARC and NMDAR complexes share 9 overlapping genes: when excluded from the analysis, we observed independent evidence for association with both gene sets (case-control-*de novo* meta-analysis: ARC *P* = 9.4 × 10^−4^; NMDAR *P* = 7.4 × 10^−5^).

**Table 2.**
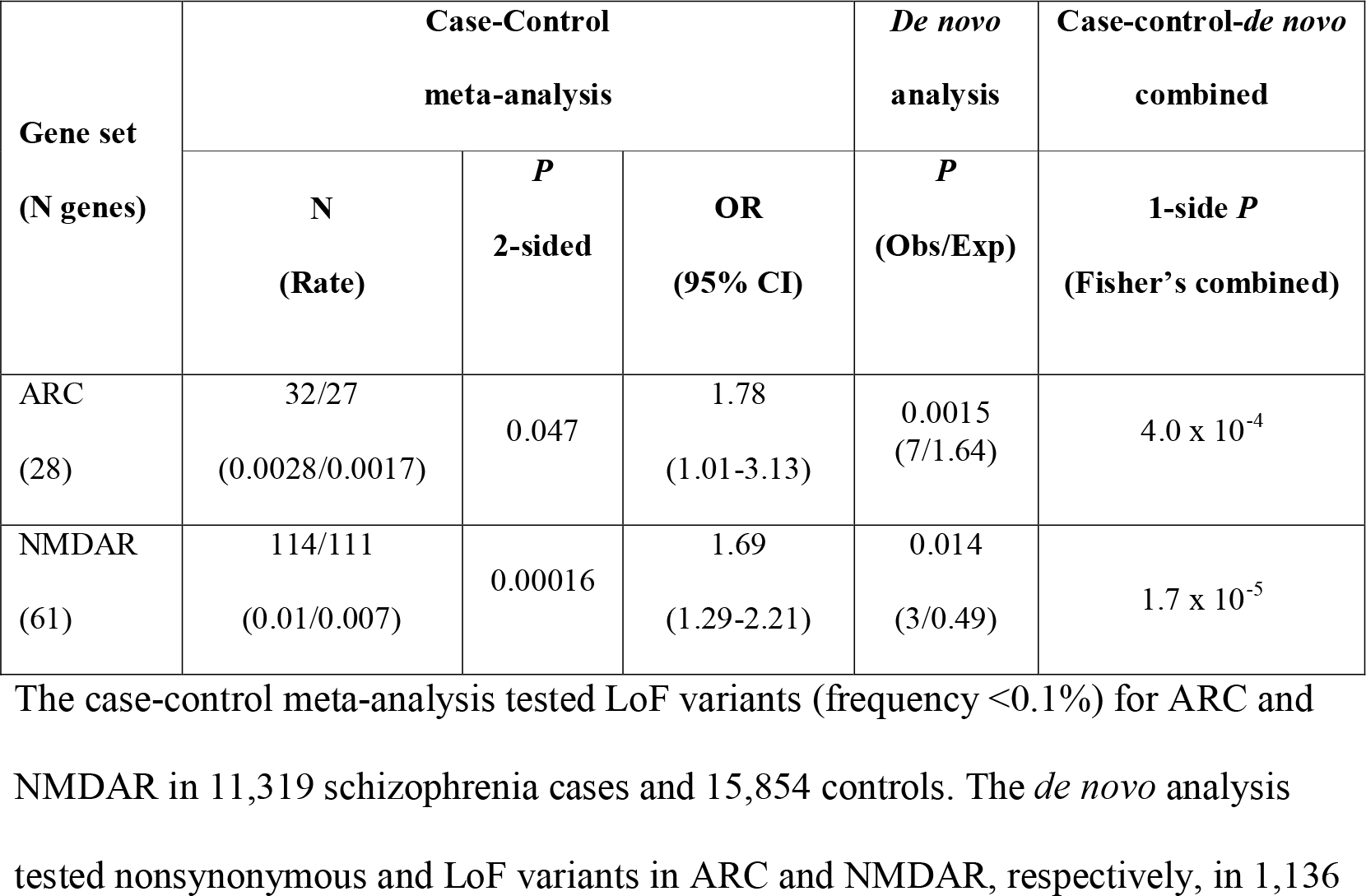
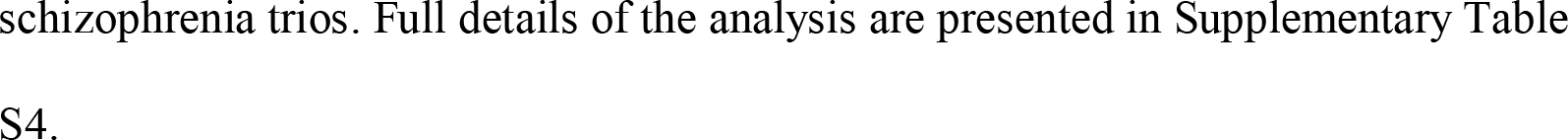
Synaptic gene set meta-analysis.

| Gene set<br>(N genes) | Case-Control<br>meta-analysis |  |  | *De novo*<br>analysis | Case-control-*de novo*<br>combined |
| --- | --- | --- | --- | --- | --- |
|  | N<br>(Rate) | *P*<br>2-sided | OR<br>(95% CI) | *P*<br>(Obs/Exp) | 1-side *P*<br>(Fisher's combined) |
| ARC<br>(28) | 32/27<br>(0.0028/0.0017) | 0.047 | 1.78<br>(1.01-3.13) | 0.0015<br>(7/1.64) | $4.0 \times 10^{-4}$ |
| NMDAR<br>(61) | 114/111<br>(0.01/0.007) | 0.00016 | 1.69<br>(1.29-2.21) | 0.014<br>(3/0.49) | $1.7 \times 10^{-5}$ |

The case-control meta-analysis tested LoF variants (frequency $< 0.1\%$ ) for ARC and NMDAR in 11,319 schizophrenia cases and 15,854 controls. The *de novo* analysis tested nonsynonymous and LoF variants in ARC and NMDAR, respectively, in 1,136
schizophrenia trios. Full details of the analysis are presented in Supplementary Table S4.

#### Voltage-gated sodium and calcium channels

We found nominally significant evidence for enrichment in cases for LoF mutations (<0.1% frequency) in voltage-gated sodium channels (targeted sequencing sample; OR (95% CI) = 1.99 (1.11-3.71); *P* = 0.02; case-control-*de novo* meta-analysis: *P* = 0.025, Supplementary Table S4), but no evidence for association between schizophrenia and voltage-gated calcium channels (Supplementary Table S4).

#### Paralog conserved ion channel sites

In the targeted sequence sample, we found a significant case excess of rare (frequency <0.1%) paralog conserved missense and LoF variants in sodium channels (OR (95% CI) = 1.26 (1.08 – 1.47); *P* = 0.0035) but not calcium channels (Supplementary Table S5). Enrichment of rare (frequency <0.1%) paralog conserved missense and LoF variants was also supported in the full case-control meta-analysis (OR (95% CI) = 1.18 (1.07 – 1.31); *P* = 0.0014, Fig 2, Supplementary Table S5), and was robust to exclusion of LoF mutations from the analysis (OR (95% CI) = 1.16 (1.04 – 1.29); *P* = 0.007). The effect size for rare (frequency <0.1%) paralog conserved missense and LoF variants was significantly different to that for paralog non-conserved missense variants (Z-test *P* = 0.0018), indeed there was no enrichment for missense variants at paralog non-conserved sites (case-control meta-analysis *P* = 0.44, Fig 2, Supplementary Table S5).

**Fig 2.**
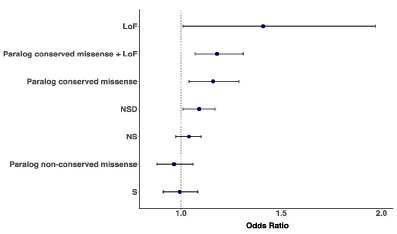
Case-control meta-analysis of rare (frequency <0.1%) variants in voltage-gated sodium channels. S = synonymous; NS = nonsynonymous; NDS = nonsynonymous damaging; LoF = loss-of-function.

We divided the voltage-gated sodium channel set into alpha (10 genes) and beta (4 genes) subunits, testing these separately; only the alpha subunits were significantly enriched for rare (frequency <0.1%) paralog conserved missense and LoF variants (Case-control meta-analysis: Alpha subunits; OR (95% CI) = 1.2 (1.08 – 1.33); *P* = 0.00086; Beta subunits, OR (95% CI) = 0.92 (0.52 – 1.62); *P* = 0.76). In all sodium channel genes, one nonsense *de novo* mutation was observed in *SCN2A* (*denovo P* value for LoF and paralog conserved missense variants in sodium channel alpha subunits = 0.75; case-control-*de novo* meta-analysis: *P* = 0.0029).

Paralog conserved analysis did not find association with schizophrenia for individual voltage-gated sodium channel genes after correction for multiple testing (Supplementary Table S6).

### Single gene analysis

In the meta-analysis (LoF; frequency <0.1%) of all data, no gene was associated with schizophrenia after Bonferroni correction (given this analysis included exome sequence data used to select our gene targets, we corrected for ~20,000 genes, Supplementary Table S7). The most significant gene was *TAF13* (*P*= 1.6 × 10^−5^), support coming mainly from published LoF *de novo* mutations as noted before [9] (Supplementary Table S7).

## Discussion

Sequencing studies have started to provide novel insights into the genetic architecture and aetiology of schizophrenia, although these are still limited by small sample sizes and low power. Seeking to increase power for a prioritized set of genes, we sequenced the coding regions of 187 schizophrenia candidates in over 10,000 samples that have not contributed to previous sequencing studies of schizophrenia.

Across all candidates, we found a significant excess of LoF alleles in the independent set of schizophrenia cases, confirming our hypothesis that one or more of the candidates is involved in schizophrenia pathogenesis. The strongest evidence for enrichment was for LoF alleles with a frequency <0.1%, suggesting that recurrent rather than only singleton schizophrenia risk alleles are present among our 187 targeted genes. This appears to contrast with a recent Swedish exome-sequencing study of schizophrenia, which reported an increased exome-wide burden in cases of ultra-rare protein altering alleles observed only once in their sample and never in 45,376 non-psychiatric ExAC individuals [7]. Our analyses (data not shown) of the same Swedish dataset confirms at an exome wide level, singleton LoF mutations to be more highly enriched than those with frequency <0.1% (after excluding singletons; Z-test *P* = 0.00035) although this did not hold when restricted to the 187 targeted genes (Z-test *P* = 0.11).

In the present study, we conducted the largest schizophrenia sequencing meta-analysis of RCVs in the synaptic gene sets ARC and NMDAR to date. The inclusion of our new independent data in this analysis strengthened the evidence for association between RCVs in ARC and NMDAR and schizophrenia. In the context of previously published research, where rare and *de novo* CNVs in these gene sets have been consistently associated with schizophrenia [5, 10, 14], the results provide a strong body of evidence for the involvement of ARC and NMDAR proteins in the aetiology of schizophrenia.

Of the other two comprehensively tested functional candidate sets, only voltage-gated sodium channels were enriched for rare (frequency <0.1%) LoF alleles in cases and this was independently supported by a novel case-control analysis of missense variants at paralog conserved sites. The same voltage-gated sodium channel gene set was previously implicated in schizophrenia in an analysis of compound heterozygous mutation [18], a model that cannot be adequately tested in the present dataset given the inability to phase very low frequency variation. In addition to previous genetic support in schizophrenia, we note sodium channels have high biological plausibility given that mutations in this gene set have been associated with other neurodevelopmental disorders, including some forms of epilepsy and developmental delay [20–22]. In the current study, no single sodium channel gene was significantly associated with schizophrenia (after correction for multiple testing), with the most significant gene being *SCN7A* (*P*_*uncorrected*_ = 0.0012).

The sodium channel set contains 14 genes, 10 encoding alpha subunits involved in generating action potentials [21], and 4 beta subunits which, in association with alpha subunits, modulate their gating and cellular excitability [22]. In the present study, the evidence for association derives from mutations in alpha subunits, although the absence of signal in beta-subunits might simply reflect low power (there are fewer beta-subunits, of which paralog conservation scores are only available for *SCN2B* and *SCN4B*, whereas paralog conservation scores are available for all 10 alpha subunits).

Despite the increased sample size, we did not observe any single-gene association that remained statistically significant after correction for multiple testing. However, our gene set enrichment results suggest that significant single gene associations will be discovered when larger samples are combined with more effective methods for identifying disease-associated missense variation.

A limitation of our study was the exclusion of indel mutations (see methods) from the targeted sequencing. Also, given the restrictions posed by targeted sequencing, we were unable to test some of the much larger gene sets that have been implicated by common and rare variation in schizophrenia, for example targets of FMRP [4, 7, 9, 23].

In conclusion, we conducted one of the largest sequencing studies of schizophrenia to date, which targeted the protein coding regions of 187 putative schizophrenia risk genes. We found a significant excess of LoF alleles in cases among all 187 targeted genes. By leveraging information from paralog conservation, we provide novel evidence that multiple voltage-gated sodium channels are involved in schizophrenia pathogenesis. We provide further support for association between RCVs in ARC and NMDAR post-synaptic protein complexes and schizophrenia. Larger samples are required to identify the specific genes and variants driving these gene set associations.

## Methods

### Sample description

#### Targeted sequence sample

A total of 11,493 blood-derived DNA samples were selected for targeted sequencing (5,724 cases and 5,769 controls). None have been included in previous schizophrenia sequencing studies. The majority of sequenced cases were from the CLOZUK dataset (n=4,647), which has been described previously [24] and in the Supplementary Material. We sequenced additional cases from the UK (Cardiff COGS cohort; n=521), Ireland (Dublin cohort; n=335) and the Netherlands (GROUP cohort [25]; n=221). The majority of sequenced controls were part of the WTCCC2 consortium (1958 birth cohort n=2,860, UK blood donors n = 2,463) [26–28]. Additional controls were sequenced from the Dublin (n= 230) and GROUP cohorts (n=216) [25]. Sample descriptions are presented in the supplementary material.

#### Additional data sets

We acquired publically available case-control exome sequencing data from two previously published schizophrenia studies, the UK10K Exome-sequencing study (N=1,352 cases and 4,769 controls) [11] and a Swedish study (N= 4,867 cases and 6,140 controls) [7]. Additionally, *de novo* mutations from 1,136 published schizophrenia-proband parent trios were derived from our own and other published whole exome sequencing studies [9, 29–36] (Supplementary Material Table S8).

### Targeted Sequencing

We designed an Ampliseq custom panel (Thermo Fisher) for targeting the exons of 187 genes. The panel comprised two pools of 3,094 and 3,082 amplicons each and covered a total region of 750kb. Library preparation used the Ion AmpliSeq protocol, using 10ng of DNA per pool, the Ion Ampliseq Kit Version 2.0 and the AmpliSeq custom panel (Thermo Fisher). We barcoded the libraries using the Ion Express Barcode Adapters 1-96 Kit (Thermo Fisher). Unamplified libraries were quantified by qPCR using Ion TQMN Quantitation kit following manufacturer’s instructions (Thermo Fisher) and diluted to 100pM. Groups of 72 uniquely barcoded libraries were combined onto single Ion Chips. Sequencing was performed on the Ion Proton benchtop sequencing platform (Thermo Fisher) following the manufacturer’s protocol. Sequencing took place in two waves, corresponding to different versions of the sequencing and chip kits: 2,305 cases and 2,274 controls were sequenced in wave 1 using the Ion PI IC 200 kit and Ion PI Chip kit v2 BC (Thermo Fisher); 3,419 cases and 3,495 controls were sequenced in wave 2 using the Ion PI HiQ kit and Ion PI Chip kit v3 BC (Thermo Fisher). Data processing and QC procedures were conducted independently for each wave. The mean target coverage for cases and controls passing QC was 158X and 160X for wave 1, and 154X and 145X for wave 2, respectively (density plots of sequence coverage in Supplementary Material Fig S1). Both cases and controls had at least 95% of target bases covered at ≥10X.

### Gene-selection

We sequenced the coding regions of genes belonging to the following gene sets: ARC (n=28) [9], NMDAR (n=61) [9], voltage-gated calcium channels (n=26) [15] and voltage-gated sodium channels (n=14) [18]. We sequenced an additional 58 genes, selected for having two or more supportive lines of evidence for association with schizophrenia (full criteria for gene-selection described in Supplementary Material). A list of all 187 sequenced genes, and the rationale for selection is presented in Supplementary Table S1.

### Data processing and quality control

Sequence data were independently processed for each Ion Torrent wave according to GATK best practice guidelines [37, 38]. Reads were aligned to the human g1k (v37) reference genome using bwa [39]. Variants were called using GATK haplotype caller (v3.4) and filtered using the GATK Variant Quality Score Recalibration (VQSR) tool.

#### Sample level QC

Individuals were excluded if they were more than 3 standard deviations from their sequencing wave’s mean for: proportion of variants in dbSNP; number of alternative alleles; number of singletons; total number of synonymous mutations; total number of nonsynonymous mutations. When available, SNP genotyping array data were used to assess sequencing-array genotype concordance (array genotypes used as truth set) and to identify duplicate/first degree relatives. SNP genotyping array data from 3 chips (Illumina OmniExpress, Illumina ExomeChip, Immunochip) were available for 96% (5,508/5,724) of cases and 72% (4,149/5,769) of controls. Samples were excluded if they had a genotype concordance < 0.9 or if they were found to be one member of a duplicate (kinship coefficient > 0.354) or first-degree relative (kinship coefficient > 0.177) pair (identified using the KING toolset [40] in the Bioconductor package SNPRelate). For samples not previously genotyped using SNP arrays, we used Ion Torrent sequence data to identify and exclude duplicate samples.

Principal component analysis (PCA) was used to identify and exclude cases and controls with non-European ancestry. We performed PCA in the 1000 genomes project data (phase 3), using variants found in both targeted sequence data and 1000 genomes data, and projected our targeted sequence samples onto these PCs using EIGENSOFT smartPCA [41]. Targeted sequence samples were excluded if PCs 1 and 2 were more than 3 standard deviations from the mean of PCs from European 1000 genome samples (Supplementary Fig S2). Post sample QC, 5,207 cases and 4,991 controls from the targeted sequence sample were retained for analysis.

#### Variant level QC

Variant sites within each targeted Ion Torrent sequencing wave were excluded if they failed Hardy-Weinberg equilibrium exact tests (χ^2^*P* < 10^−8^), GATK VQSR filters, had > 20% missingness, or contained an indel (sequence data produced by Ion Torrent instruments has low indel specificity [42]). For targeted sequencing data, individual genotypes were set to missing if they had a depth (DP) < 20, genotype quality (GQ) < 80, allele-balance (AB) < 0.9 for non-reference homozygous genotypes or an AB < 0.2 or > 0.8 for heterozygous genotypes. For analysis of previously published exome sequencing data, we applied filters which more closely matched those described in their original publications (DP <=10, GQ < 30, AB < 0.9 for non-reference homozygous genotypes, AB < 0.2 or > 0.8 for heterozygous genotypes) [7, 11]. All variant filtering was conducted using Hail software (https://github.com/hail-is/hail) [43].

### Variant annotation

In primary burden tests, we analysed three classes of mutation (LoF, nonsynonymous damaging, and nonsynonymous) and two allele frequency thresholds (<0.1% and singletons). We defined LoF variants as those producing premature stop codons (nonsense) or situated at essential splice sites (within 2 bases either side of exon junctions). Damaging nonsynonymous mutations were defined as LoF alleles and missense alleles with PHRED-scaled CADD score ≥ 20 (representing the predicted top 1% most deleterious variants in the genome) [44]. For analysis of previously published case-control data (UK10K and Swedish samples), which were exome sequenced on Illumina instruments, we included frameshift indels as LoF mutations and frameshift/in-frame indels as nonsynonymous mutations. We annotated and filtered variants using the frequencies observed in their respective data set (Ion Torrent wave 1, Ion Torrent wave 2, UK10K or Swedish) and each ExAC sub-population (European (Non-Finnish), African, East Asian, European (Finnish), Latino, Other, South Asian) [16]. Singletons were annotated as alleles observed once in all available sequence data (targeted, UK10K and Swedish data) and never in 45,376 individuals without a known psychiatric diagnosis from the Exome Aggregation Consortium [16].

To annotate sites with their paralog conservation scores, we used para_zscores downloaded from https://zenodo.org/record/817898. In our paralog conserved analysis, we followed the publication describing this metric [20] by testing the burden of all LoF alleles and missense alleles at sites annotated as having a para_zscore > 0.

Variants were annotated using Hail’s ensemble VEP method (version 86, http://oct2016.archive.ensembl.org/index.html).

### Statistics

**Case-control analysis:** Gene set and single gene association statistics were generated using the following Firth’s penalised-likelihood logistic regression model.

Logit (pr(case)) ~ N test variants + baseline synonymous count + first 10 PCs + sex + Ion Torrent sequencing wave (targeted analysis only).

N test variants refers to the number of putative risk alleles observed in each sample (e.g. number of LoF singletons in the gene/gene set tested). Baseline synonymous count refers to the number synonymous alleles observed in all sequenced genes, using the same allele frequency threshold used for the test variant (e.g., if LoF singletons are the test variant, then the overall number of synonymous singletons are corrected for). This covariate was only included in tests of nonsynonymous or LoF mutation to control for potential unknown technical biases [7].

Test statistics were generated independently for each case-control dataset (targeted, UK10K and Swedish), using the logistf function implemented in R (version 3.3.1). Odds ratios (ORs) for the increased risk of schizophrenia incurred for each mutation were obtained from the Firth’s penalised-likelihood logistic regression model described above.

***De novo* mutation analysis:** Enrichment of *de novo* mutation in genes/gene sets wastested using the statistical framework described in Samocha *et al* 2014 [45]. Here, gene mutation rates provided in Ware *et al* 2015 [46] were used to estimate the expected number of *de novo* mutations in the gene/gene set, which was then compared to the observed number of *de novo* mutations using a Poisson test (implemented in R). As gene mutation rates for in-frame indels are not provided in [46], we adopted the method used by the Deciphering Developmental Disorders Study [47], which scaled frameshift mutation rates by the ratio of in-frame to frameshift mutations reported to occur in genome-wide regions under weak negative selection (ratio 1:9) [47]. To estimate the expected number of LoF *de novo* mutations and missense *de novo* mutations at paralog conserved sites in sodium channel alpha subunits, we used the mutation rates provided in [20].

**Meta-analysis:** Coefficients and standard errors from independently analysed case-control (targeted, UK10K and Swedish) regression tests were meta-analysed as fixed effects using the inverse-variance method (implemented in R using the rma.uni() function as part of the metafor package). To obtain a single enrichment statistic for meta-analysed case-control and *de novo* tests, we followed the method described in [11], which combined a 1-tail case-control *P* value with the *de novo* Poisson test P value using Fisher’s combined method. For our combined case-control-*de novo* meta-analysis of nonsynonymous damaging mutations, we included all *de novo* nonsynonymous mutations (i.e. not just those with a CADD score ≥ 20), given they are *a priori* more likely to be deleterious than inherited variation [48] and were the class of mutation most strongly associated with schizophrenia candidate genes in our previous publication [9].

## Acknowledgments

The work at Cardiff University was primarily funded by Medical Research Council (MRC) Centre (MR/L010305/1) and Program (G0800509) Grants Additional support was provided from the European Community’s Seventh Framework Programme HEALTH-F2-2010-241909 (Project EU-GEI) and the European Union’s Seventh Framework Programme for research, technological development and demonstration under grant agreement n° 279227 (CRESTAR Consortium).

We thank the participants and clinicians who took part in the CardiffCOGS study. For the CLOZUK2 sample we thank Leyden Delta for supporting the sample collection, anonymisation and data preparation (particularly Marinka Helthuis, John Jansen, Karel Jollie and Anouschka Colson), Magna Laboratories, UK (Andy Walker) and, for CLOZUK1, Novartis and The Doctor’s Laboratory staff for their guidance and cooperation.

We acknowledge Kiran Mantripragada, Lesley Bates and Lucinda Hopkins, at Cardiff University, for laboratory sample management. We acknowledge Wayne Lawrence and Mark Einon, at Cardiff University, for support with the use and setup of computational infrastructures. We acknowledge Tarjinder Singh, Jeffrey Barrett, and other members of the UK10K consortium (https://www.uk10k.org/) [11].

## GROUP acknowledgments

GROUP investigators are: Behrooz Z. Alizadeh^a^, Therese van Amelsvoort^g^, Agna A. Bartels-Velthuis^a^, Nico J. van Beveren^b,c,d^, Richard Bruggeman^a^, Wiepke Cahn^e^, Lieuwe de Haan^f^, Philippe Delespaul^g^, Carin J. Meijer^f^, Inez Myin-Germeys^h^, Rene S. Kahn^e^, Frederike Schirmbeck^f^, Claudia J.P. Simons^g,i^, Neeltje E. van Haren^e^, Jim van Os^g,j^, Ruud van Winkel^g,h^, Jurjen J. Luykx^c,k,l,m^

^a^ University of Groningen, University Medical Center Groningen, University Center for Psychiatry, Groningen, The Netherlands;

^b^ Antes Center for Mental Health Care, Rotterdam, The Netherlands;

^c^ Erasmus MC, Department of Psychiatry, Rotterdam, The Netherlands;

^d^ Erasmus MC, Department of Neuroscience, Rotterdam, The Netherlands;

^e^ University Medical Center Utrecht, Department of Psychiatry, Brain Centre Rudolf Magnus, Utrecht, The Netherlands;

^f^ Academic Medical Center, University of Amsterdam, Department of Psychiatry, Amsterdam, The Netherlands;

^g^ Maastricht University Medical Center, Department of Psychiatry and Psychology, School for Mental Health and Neuroscience, Maastricht, The Netherlands;

^h^ KU Leuven, Department of Neuroscience, Research Group Psychiatry, Center for Contextual Psychiatry, Leuven, Belgium;

^i^ GGzE, Institute for Mental Health Care Eindhoven and De Kempen, Eindhoven, The Netherlands;

^j^ King’s College London, King’s Health Partners, Department of Psychosis Studies, Institute of Psychiatry, London, United Kingdom;

^k^ University Medical Center Utrecht, Department of Translational Neuroscience, Brain Centre Rudolf Magnus, Utrecht, The Netherlands;

^l^Department of Psychiatry, ZNA Hospitals, Antwerp, Belgium;

^m^SymforaMeander hospital, medical-psychiatric unit, Amersfoort, the Netherlands

The infrastructure for the GROUP study is funded through the Geestkracht programme of the Dutch Health Research Council (Zon-Mw, grant number 10-000-1001), and matching funds from participating pharmaceutical companies (Lundbeck, AstraZeneca, Eli Lilly, Janssen Cilag) and universities and mental health care organizations (Amsterdam: Academic Psychiatric Centre of the Academic Medical Center and the mental health institutions: GGZ Ingeest, Arkin, Dijk en Duin, GGZ Rivierduinen, Erasmus Medical Centre, GGZ Noord Holland Noord. Groningen: University Medical Center Groningen and the mental health institutions: Lentis, GGZ Friesland, GGZ Drenthe, Dimence, Mediant, GGNet Warnsveld, Yulius Dordrecht and Parnassia psycho-medical center The Hague. Maastricht: Maastricht University Medical Centre and the mental health institutions: GGZ Eindhoven en De Kempen, GGZ Breburg, GGZ Oost-Brabant, Vincent van Gogh voor Geestelijke Gezondheid, Mondriaan, Virenze riagg, Zuyderland GGZ, MET ggz, Universitair Centrum Sint-Jozef Kortenberg, CAPRI University of Antwerp, PC Ziekeren Sint-Truiden, PZ Sancta Maria Sint-Truiden, GGZ Overpelt, OPZ Rekem. Utrecht: University Medical Center Utrecht and the mental health institutions Altrecht, GGZ Centraal and Delta.)

We are grateful for the generosity of time and effort by the patients, their families and healthy subjects. Furthermore we would like to thank all research personnel involved in the GROUP project, in particular: Joyce van Baaren, Erwin Veermans, Ger Driessen, Truda Driesen, Karin Pos, Erna van ‘t Hag, Jessica de Nijs, Atiqul Islam, Wendy Beuken and Debora Op ‘t Eijnde.

## Supporting Information Captions

**Table S1**. Genes targeted for sequencing. Gene IDs are presented for all 187 targeted genes, along with the criteria used to select them for sequencing.

**Table S2.** Targeted sequence sample case-control LoF mutations. All loss of function (LoF) mutations observed in new targeted sequence data, for alleles < 0.1% in frequency. Singletons are indicated in the Is_Singleton column.

**Table S3.** Total-burden analysis. Case-control association results for all 187 targeted genes and 106 LoF intolerant genes (genes with pLi scores > 0.9). Results for variants with a frequency < 0.1% are shown in tab 1, and for singleton variants in tab 2.

**Table S4.** Primary gene-set analysis. Gene set association results for all three case-control datasets (Targeted sequence sample, Swedish, UK10K) and case-control-*denovo* meta-analysis (Fisher’s combined method). Results for variants with a frequency < 0.1% are shown in tab 1, and for singleton variants in tab 2.

**Table S5.** Ion Channel gene set analysis of LoF and paralog conserved missense variants (<0.1% frequency). Paralog conservation scores (para_zscores) were downloaded from https://zenodo.org/record/817898.

**Table S6.** Single-gene meta-analysis of sodium channel genes for LoF and paralog conserved missense variants (<0.1% frequency).

**Table S7.** Primary single-gene meta-analysis of LoF variants (<0.1% frequency). Single-gene results for all three case-control (Targeted sequencing, Swedish, UK10K) and *de novo* mutations.

